# Rusty Blackbirds *(Euphagus carolinus)* appear to exhibit predator satiation with a resident raptor species

**DOI:** 10.64898/2026.07.09.737236

**Authors:** Xavier R. Martin, Samuel R. Ward, Brian C. Hibbard, Oscar Gonzalez

## Abstract

Predator satiation is an understudied topic among vertebrate organisms, especially avian species. In Anderson, South Carolina, a species of migratory black bird, *Euphagus carolinus* (Rusty Blackbird) was observed roosting seasonally in a patch of *Phyllostachys aurea* (Golden bamboo) in large numbers. A resident avian predator, *Buteo lineatus* (Red-shouldered Hawk was seen preying on these roosting blackbirds; however, when this hawk would attack, pin down, and eat a blackbird, the rest of the flock would not react or fly away. Upon initial observation, we questioned if these roosting blackbirds exhibited predator satiation as a protection mechanism against avian predation. When *E. carolinus* flocked to roost, their numbers were visually counted and compared between at least two members of the research team. After allowing the birds to settle between fifteen seconds and one minute, a *B. lineatus* call was played with speakers to the flock. Simultaneously, a regulation-sized football was tossed to simulate the hawk attacking a member of the flock. The number of birds that flew away was recorded. Results between roosting numbers and flight numbers upon disturbance were analyzed to see if roosting group size correlates with evasiveness in birds. Results indicated that as blackbird group size increased, the total percentage of escaping birds decreased. Statistical analysis showed the leaving percentage versus the natural logarithm of group size was highly statistically significant (*p* < 0.001) and had a moderately negative overall correlation (*rs* = -0.62). Our results suggest that these blackbirds feel safer in large groups instead of small groups during predation events when roosting in dense brush.

**LAY SUMMARY:**

- There is little research involving predator satiation tendencies in birds.
- Predator satiation is a strategy developed by prey animals to cope with predators, relying on sheer numbers to avoid predation instead of fight or flight.
- We observed roosting Rusty Blackbirds in Anderson, SC displaying predator satiation-linked tendencies under avian predation stress when in large groups.
- We tested the behavior of Rusty Blackbirds by simulating a hawk attack, noting the group size and the number of birds escaping.
- Our results showed that as group size increased, flight percentage decreased under a simulated predator attack.
- Rusty Blackbirds seem to use the predator satiation strategy when roosting.

## INTRODUCTION

The role of predators is crucial in maintaining the balance of an ecosystem. Though most avian species have evolved unique physiological tendencies to ward off or evade potential predators, many migratory bird species have not, as new terrain increases predation hazard during migratory stopovers (Swaddle 1998, Dierschke 2003). Survival rates increase when avian prey adapts to a predator’s tendencies or avoids the predator altogether. The impact of predation extends beyond individual species; it can shape entire ecosystems (Glasser 1979). The decline or disappearance of a predator can result in the overpopulation of certain prey species, degrading habitat and reducing biodiversity over time. Conversely, a healthy predator population promotes balance, encouraging diverse communities of plants and animals. Predators can drive populations to adapt to their feeding tendencies uniquely, through camouflage, mimicry, or behavioral changes like flocking or flying evasively.

One seemingly bizarre method that increases prey survival is predator satiation. This strategy involves prey species overwhelming their predators by producing many offspring or migrating in large groups, increasing individual survival probability. Prey species reliant on predator satiation allow predators to gorge until satiated; however, the sheer number of prey will be far too great for predators to make more than a dent in the total population. Predator satiation is often seen in R-selected species such as cicadas, mayflies, and baitfish. Most R-selected species exhibiting predator satiation are physically inferior to their pursuers, sacrificing defensive prowess for population density (Neill and Cullen 1974, Sweeney and Vannote 1982, Williams et al. 1993). Species such as cicadas have been observed to rely on predator satiation, as they show little fear or interest in predators, relying on sheer numbers for protection. Cicadas emerge in great multitudes, overwhelming predators, who become exhausted from eating them, allowing most of the cicadas to thrive (Karban 1982, Machta et al. 2019). Predator satiation has been researched mainly in invertebrate and plant species; the research linked to vertebrate species is limited at best (Greenberg and Zarnoch 2018, Zwolak et al. 2022).

We suspected predator satiation as a strategy used by *Euphagus carolinus* (Rusty Blackbirds) in South Carolina, United States. Rusty Blackbirds are a species of migratory songbird that breed in the northern forests of Alaska and Canada (Wohner et al. 2020) and are usually observed as black with hints of brown over their crown to the under-tail coverts: a color warranting the “rusty” part of their name (Avery 2020). *E. carolinus* have elliptical wings, making them exceptionally effective at flying quickly in short bursts through the forests they inhabit. Anatomically, elliptical wings are found on granivorous and insectivorous bird species, proving useful for insect capture and predator evasion (van den Hout et al. 2009). Their migration begins in September and continues in the southeastern United States throughout the winter. During the fall migration, these birds feed primarily on berries, seeds, and insects in open fields and swamps. They prefer to roost in high, dense thickets near wetland habitats with many other blackbird species and *Quiscalus quiscula* (Common Grackles) but typically forage in single-species flocks (Greenberg et al. 2011).

Large populations of *E. carolinus* can be observed all over Anderson, South Carolina during seasonal migration (Figure 1). *Buteo lineatus* (Red-Shouldered Hawks) also lived nearby and were observed attacking roosted *E. carolinus* in the fall of 2024. The hawk grabbed a roosting *E. carolinus*, killed it, and flew away with its prey. There were also several (1-5) dead blackbirds on the ground at any point during peak migration, with organs ripped out, heads missing, and flesh visible, which are typical signs of predation. It has been observed that *E. carolinus* continue to roost around their fallen flock members and will pay little to no attention to predation events. However, when a solo bird or small group of birds landed, they startled easily compared to large groups. These are rare observations because *B. lineatus* do not typically eat blackbirds. Disturbance or predation also usually trigger fight or flight behaviors in blackbirds but did not in this case (Rodriguez-Prieto et al. 2009, Strobel and Boal 2010).

**Figure 1.**
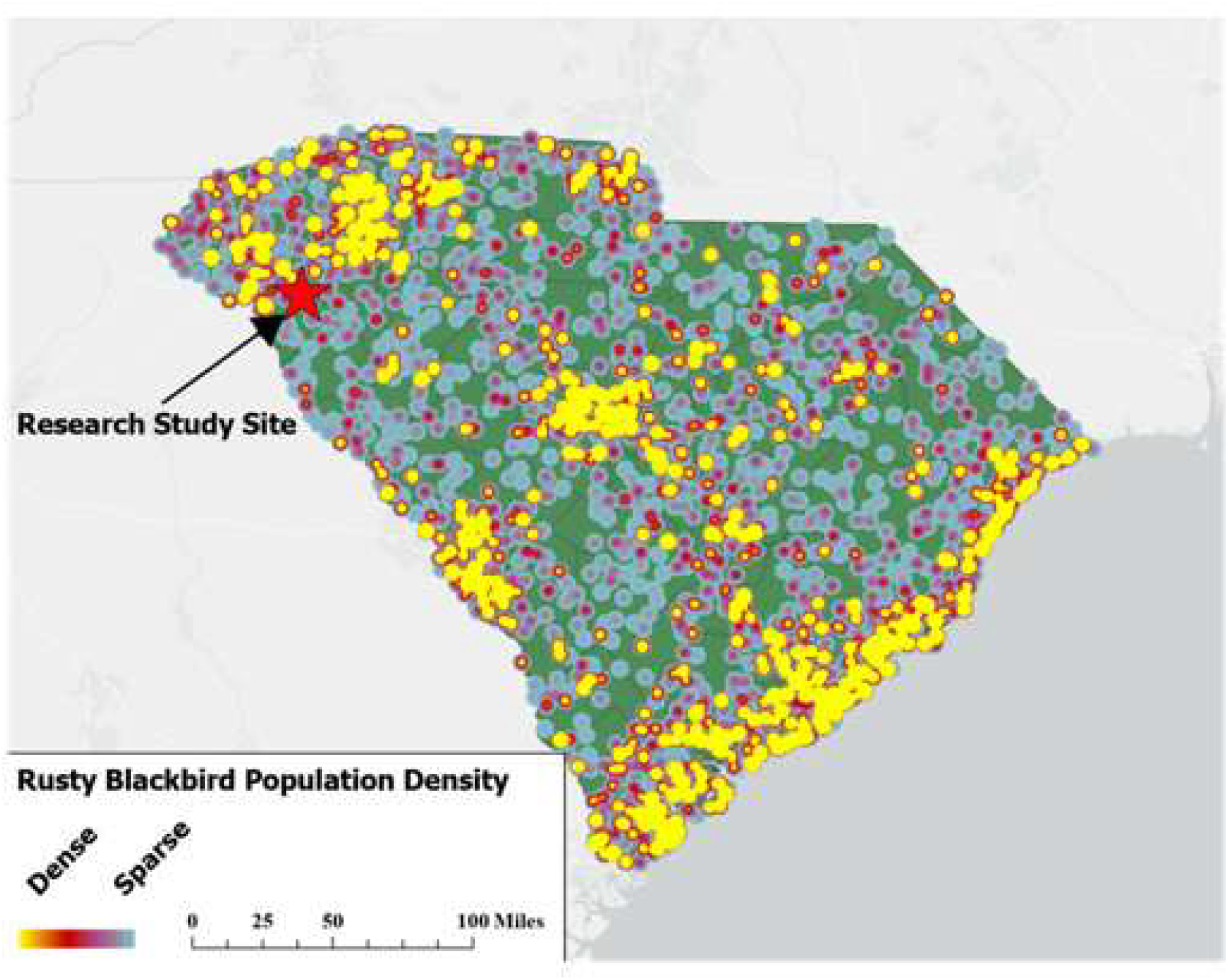
Rusty Blackbird Population data in South Carolina 2014-2024. There are high concentrations of these birds in three major areas: upstate, central, and coastal (Fink et al. 2024).

Upon observation, we hypothesized that as *E. carolinus* group numbers increased, they began to rely on predator satiation for survival instead of predator evasion. If *E. carolinus* were relying on predator satiation, we predicted that as their roosting group size increased, flight response to a predation simulation would decrease. In theory, large groups would react at a lower rate upon predation simulation, while solitary birds or small groups would fly away at a higher rate.

## METHODS

A thick patch of *Phyllostachys aurea* (Golden bamboo) next to Anderson University’s disc golf course (34.51° N, 82.63° W) was a roosting spot for over 1,000 birds every night during peak migration (September 15th through October 20th). This patch was around 61 m long by 12 m wide, making it small enough to observe in its entirety (Figure 2). Our study site was near multiple open fields, wetland areas, and *Cornus spp*. (Dogwood) habitat, making this location ideal for grain and insect forage. The migratory patterns and foraging habits were particularly evident as large migratory groups of *E. carolinus* sought sustenance and shelter in the biodiverse environment surrounding the bamboo grove.

**Figure 2.**
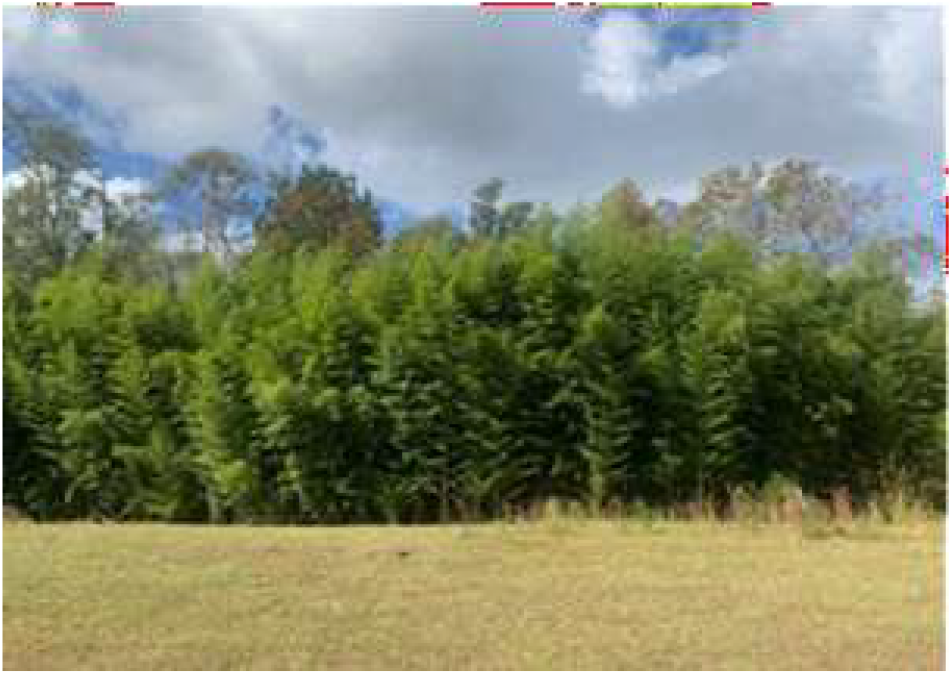
Bamboo patch in Anderson, South Carolina used as roosting site for Rusty Blackbirds. This place was used as our simulation zone.

For each predation event, two researchers counted blackbirds landing to roost on either side of the bamboo patch. The third researcher observed the number of *E. carolinus* evading the predation simulation by standing 30 m away from the bamboo patch, observing the area in its entirety. Before a trial was completed, the roosting group size was estimated based upon the following criteria: the number of individuals observed flying into or out of the bamboo patch, those heard within the patch, and the estimated number of individuals present in the bamboo patch prior to the first trial.

A total of 48 trials were conducted during October through November of 2024 and September through November 2025 to find any correlation between group size evasion percentage. The group size was log-transformed (natural log) for the analysis. We analyzed the results with the Spearman rank correlation coefficient (*rs*), and the coefficient of determination (*r*^2^) to identify the significance of the results.

## RESULTS

During the first season of research (2024), group sizes ranged from 1 to 20 blackbirds with 34 trials conducted. During the second season of research (2025), group sizes ranged from 2 to 500 blackbirds with 14 trials conducted. A total of 2,715 *E carolinus* were recorded during this study. Roosting group size and evasive group size were recorded after every trial. The outcomes of the two years’ trials were examined both in isolation and in direct comparison with one another (Table 1).

**Table 1.** Summary statistics of the predation simulations with *Euphagus carolinus*; comparing the roosting group size with the birds that left. Spearman’s rank correlation coefficient (*rs*), coefficient of determination (*r*^2^) and p-value for each year’s analysis. Mean (Average group size) and standard deviation (σ) are also included.

| Analysis | Year 1 (2024) | Year 2 (2025) | Combined results |
| --- | --- | --- | --- |
| $rs$ | -0.6506 | -0.4842 | -0.6218 |
| $r^2$ | 0.4233 | 0.2345 | 0.3867 |
| p-value %Leaving<br>birds vs. ln(Group<br>size) | <0.001 | 0.1562 | <0.001 |
| Average group size | 5 | 182 | 57 |
| $\sigma$ | 5 | 171 | 121 |

The first year of results indicated that the percentage of birds escaping declined as the resting bird group size increased. A line of best fit indicated a steep decline in the percentage of escaping birds until ∼10 birds, when it began to decelerate to around 10% (Figure 3A). Statistical analysis indicated that Spearman’s correlation coefficient had a moderately negative relationship with group size and the percentage that displayed evasion tactics during predation (*rs* = -0.65).

**Figure 3.**
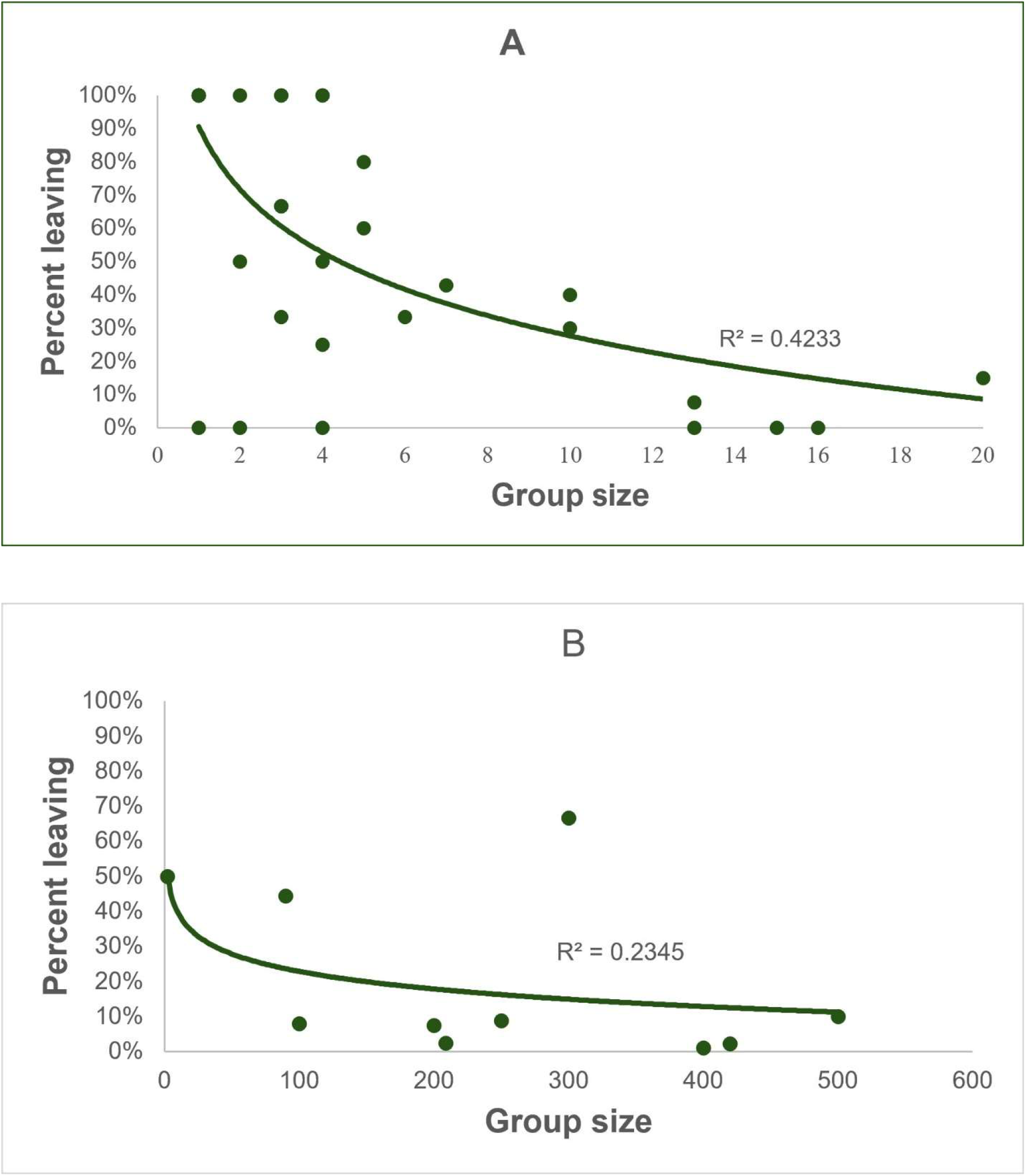
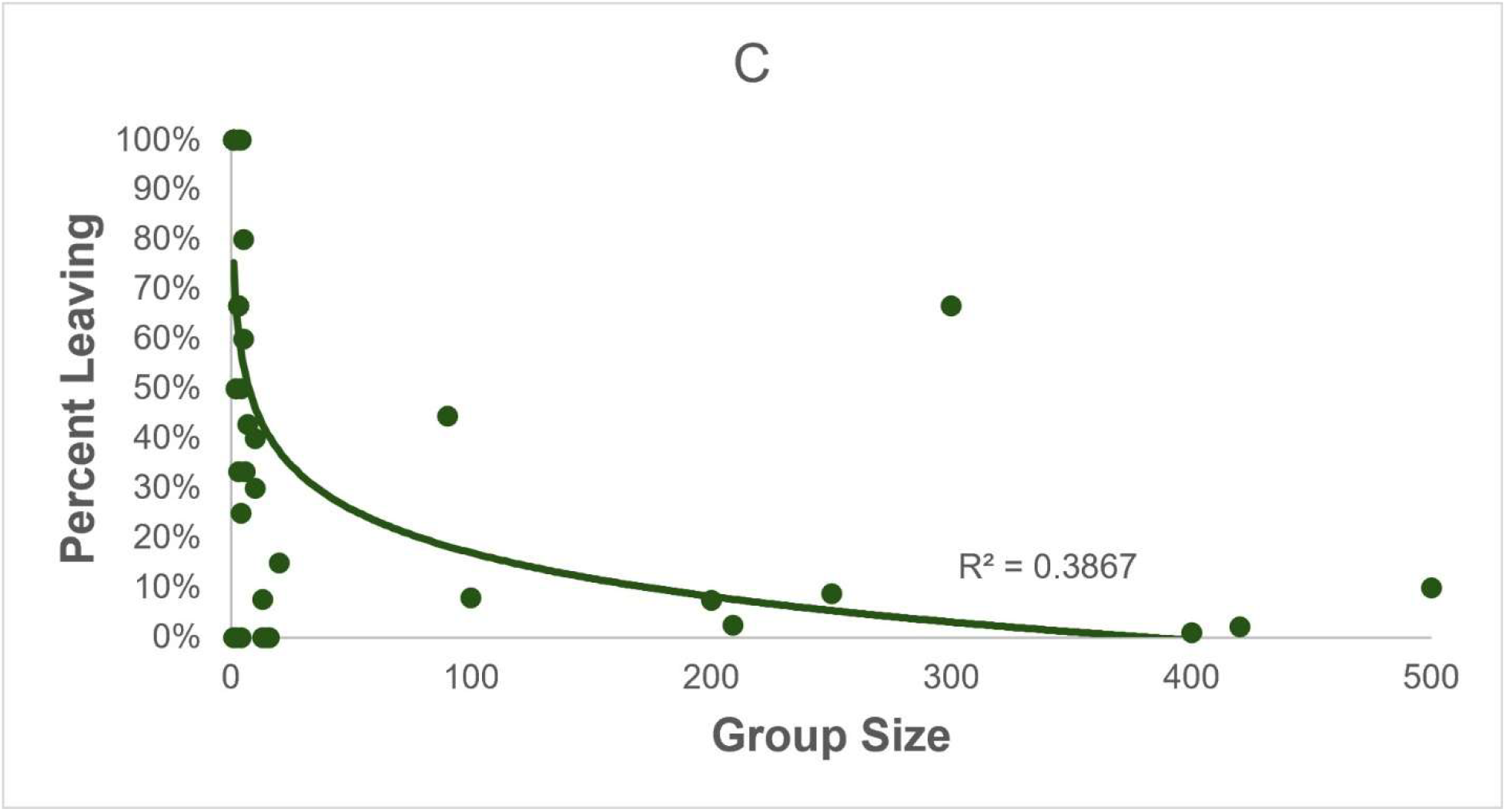
Results of predation simulation on roosting *E. carolinus*. The X-axis is the size of the roosting group and the Y-axis is the percentage of birds that escaped after the simulation. A. Results in 2024 (*n*= 34 trials). B. Results in 2025 (*n*= 14 trials). C. Results in 2024-2025 (*n*= 48 trials).

This result suggests that larger groups tended to have a lower percentage of birds that left the group when a predation event occurred. Building upon this, the coefficient of determination value (*r*^2^ = 0.42) explained 42% of the variation in the percentage of birds leaving could be explained through the data. However, the logarithmic regression (y = -0.274ln(x) + 0.907) indicates the data was highly statistically significant when comparing the relationship of the percentage of birds that left against ln(group size) (*p* < 0.001).

Much larger groups came with the second year of results, as tests were run earlier in the year during peak migration. These results showed a much more random distribution of percentages of blackbirds escaping by group size. Nonetheless, there continues to be a relative downward trend with a decline until ∼50 birds, where it then begins to level out (Figure 3B). While not as strong as the first set of trials, statistics showed a less-than-moderate, negative relationship between group size and the percentage of blackbirds that left (*rs* = -0.48). Similarly, the coefficient of determination (*r*^2^ = 0.24) showed that 24% of the variance was explained through our data. With the calculated logarithmic relationship p-value (*p* = 0.16), the second year of data would not be statistically significant and did not have an independently robust relationship in this sample when isolated from year one.

When the results from both years were compared, highly significant results were gathered (Figure 3C). With a correlation coefficient of *rs* = -0.62, the results yielded a moderate relationship. Furthermore, the calculated coefficient of determination value (*r*^2^ = 0.39) explains 39% of the variation. Most notably, the calculated p-value of leaving percentage versus ln(group size), indicated high statistical significance (*p* < 0.001) as shown in Table 1.

## DISCUSSION

Results supported our hypothesis, showing a statistically significant negative correlation between flock size and evasion percentage. Year one group size results had the tendency to be smaller while group sizes in the second year were much larger. When comparing the data from both years, most of the groups that were larger than 10 had an evasive rate that was ≤ 10%. This could potentially suggest *E. carolinus* deem these groups sizes give them the highest probability of survival when a predation event occurs.

Similar scenarios have been identified in flocks of the marine waterfowl species, *Tringa tetanus* (Redshanks). These birds take advantage of a phenomenon known as the dilution effect. The idea of this effect is that larger groups have a lower chance of being victim to a potential predator. A single bird, separated from a group, has a high chance of predation; on the other hand, in groups larger than one, the probability of survival increases, as the predation rate is dispersed across the flock (Cresswell 1994). Though these birds exhibit this phenomenon to increase their effectiveness while foraging, the principle can still be applied in the *E. carolinus* scenario. Seasonal preparation may be a reason for the lack of evasive tactics exhibited in this group of blackbirds. It has been suggested that in preparation for winter, blackbirds will increase their energy store to decrease their starvation risk. *E. carolinus* may accept the higher predation risk resulting from increased mass as a necessary tradeoff to prevent starvation, utilizing predator satiation as a potential defense strategy (Cresswell 2002).

Within the trials, there existed several outliers that deviated the data abnormally and caused the interpretation of the results to be skewed. One value to note from the results of both years combined is the x-intercept. The logarithmic model for the percent of birds leaving versus the group size has an x-intercept of approximately 379 (Figure 3C). At this point, when a predation event occurs at 379 individuals, the graph predicts none of the blackbirds will flee. The issue lies with groups larger than 379; however, the data would then predict blackbirds will then begin to fly to the predator rather than away from it. While this is something to note with the data, it is important to note that other outliers in the data, or not enough large groups in the data, could be the reason for this oddity in the graph. Another influential piece of data would be a result from the second year of testing (See Figure 3B, group size = 300; evasive group size = 200; percent leaving = 66.7%). Removing this trial from the second-year data alters the p-value to make this year’s data statistically significant (*p* = 0.007). This trial could suggest there was another factor that caused a higher-than-average rate of birds to fly away during the simulation (i.e. other nearby college students, whistles, cars). Similarly, the influence on the first trial of the second year has a bigger impact on the p-value (group size = 2; evasive group size = 1; percent leaving = 50.0%). Removing this specific test from the data, shows the importance of smaller group sizes to the p-value (*p* = 0.55). The correlation coefficient is also affected by data outside of the trend as well. Removing instances in the first year where the percentage leaving was equal to zero causes the correlation coefficient to increase in strength and p-value to become statistically significant (*rs* = -0.79; *p* < 0.001). These could have been a result of human error as the simulation could have been conducted close enough for the blackbirds to feel threatened about a potential predation event from the ground.

Another factor to consider in this study is that *B. lineatus* does not commonly prey upon adult birds. Multiple studies across North America have noted that birds are a small fraction of this hawk’s diet, and these are typically nestlings (Portnoy and Dodge 1979, Penak 1982, Bednaz and Dinsmore 1985). It is improbable that *E. carolinus* would develop a predator satiation strategy for this infrequent predator; however, our findings still indicate a safety-in-numbers behavior may be present. *E. carolinus* are also more neophobic toward novel objects in feeding grounds compared to other blackbird species (Greenberg and Mettke-Hoffmann 2001). This raises questions about the birds’ flighty behavior and possible differences in defensive behaviors between roosting and feeding birds. General neophobic behavior toward novel objects could explain why some large groups of birds flew away in our simulations. If the football was thrown too high and became fully visible to a group of *E. carolinus*, their fear of the new object could have spurred the large group to flee. Because *E. carolinus* are typically flighty, and *B. lineatus* are infrequent predators, the escape responses likely reflect generalized antipredator vigilance instead of a specific defense against hawks. The fact that smaller groups were more likely to fly away supports this notion because they have more need to be wary with fewer eyes around to assess threats.

Future research would entail the use of infrared cameras, potentially on a drone if possible. In this way, movements in the dense foliage would be better seen, and group numbers could be exact instead of visually estimated. Researchers would be able to watch birds react in real time to predation events using aerial footage. A drone would allow researchers to observe the entire area and decrease the effect of human error due to poor visibility. Increasing the number of trials, without causing a change in the birds’ natural behaviors, would also strengthen the data and account for potential outliers.

Our work would provide insightful information into the predator-prey dynamics of prey bird species, like *E. carolinus*, and predatory species of birds, like *B. lineatus*. Studying these relationships allows the research community to have a better understanding of the ecological dynamics within our environment. Predator satiation is unique among these avian species; because the vulnerability to predators between the nesting and post-fledging periods may lead to differences in habitat selection between these stages of the breeding season (Wohner et al. 2020). Continuing this research could be vital in conserving the Rusty Blackbird population in the future. Populations of blackbirds are in a sharp decline, with up to 78% loss in the last decade, due to the loss of their boreal habitats in Canada, increased forestry services in their breeding grounds causing nests to be built in less-than-ideal habitats (Wilson et al. 2021). These poor nesting situations cause these blackbirds to have an increase in nestling mortality (Luepold et al. 2015). These birds travel to great lengths to reach specific wintering and breeding conditions, and these conditions are critical for birds to rest and refuel for their continued migration further south, or back to the north (Wright et al. 2020). Monitoring their population, behaviors, and interactions within their environment is vital in maintaining their population at sustainable levels.

## ACKNOWLEDGMENTS

We thank the students at Anderson University and Universidad Peruana Cayetano Heredia for providing insightful questions about our research, as well as Dr. Todd Fenstermacher of Anderson University for providing insight on statistical analysis. This research was approved by the Animal Welfare and Usage Committee (AWUC) of Anderson University

